# Person knowledge is independently encoded by allocentric and egocentric reference frames within separate brain systems

**DOI:** 10.1101/2023.09.08.556952

**Authors:** Robert S. Chavez, Taylor D. Guthrie, Jack M. Kapustka

## Abstract

Humans spontaneously infer information about others to perceive the similarity in behaviors and traits among people in our social environments. However, evaluating the similarity among people can be achieved through multiple frames of reference: other-to-other differences (allocentric similarity) or self-to-other differences (egocentric similarity), which are difficult to dissociate with behavioral measures alone. Here, we use functional magnetic resonance imaging to test whether the similarity of brain response patterns when thinking of others and the self is predicted by models of allocentric and egocentric similarity in the trait-judgments of well-acquainted peers from 20 independent groups of people (total N = 108; within-subject design). Results show that both allocentric and egocentric similarity during person representation are reflected in brain response similarity patterns when thinking of others but do so differentially and in non-overlapping brain systems. Specifically, allocentric similarity was positively related to brain similarity patterns in distributed regions involved in social cognition as well as in regions involved in general non-social spatial reference frame encoding. Egocentric similarity was inversely related to brain response similarity but only within dorsal regions of the medial prefrontal cortex and anterior cingulate, suggesting that activation within this region is important for dissociating self-representation from that of similar others. These results suggest that the brain processes both kinds of reference frames independently to encode trait information about conspecifics that we use to represent person knowledge about other people within realworld social networks.

---

Humans live, work, and play among people with a variety of dispositions and reputations within our social environments. Understanding the traits, behaviors, and individual characteristics of other people—a process of generating what is called person knowledge (Anzellotti Young, 2020)—within our social networks is important for establishing friendships (Byrne, 1969), reducing conflict and uncertainty (Kets Sandroni, 2019), and promoting cooperation (Schroeder et al., 2015) and collaboration towards shared goals. Despite relying on a variety of high-level information processing, human social cognition is fast and spontaneous. Within seconds (Ambady Rosenthal, 1993), people can automatically ascribe trait characteristics to others that are highly correlated with independent judgements at longer timescales and predict real-world outcomes (Todorov et al., 2005) without explicit impression formation goals (Winter Uleman, 1984). These capacities form the basis of our ability to encode person knowledge that we use to maintain lasting impressions of the people with whom we interact regularly and have the potential to significantly impact our daily lives.

When using person knowledge to represent information about other people in social contexts, it is useful to know the similarity among individuals within a group. For instance, if two coworkers share many of the same traits or characteristics (e.g., both are quiet but hard working), it may be cognitively efficient to mentally represent information about them more similarly than we would if they did not share these characteristics (Macrae et al., 1994). Likewise, if we perceive someone as being similar to ourselves, we will anchor on our own attitudes when considering the preferences of others (Tamir Mitchell, 2010) which simplifies elements of social judgment and decision making (Tamir Mitchell, 2012). Thus, previous research indicates that we use information about a person’s similarity to others, as well as their similarity to ourselves, when we are generating a mental representation of their traits and behaviors. However, it is unclear whether these two ways of representing person knowledge are processed independently of one another or if they are just different operations of the same social cognitive system.

To better understand whether these two processes represent fundamentally distinct models of how we represent information about others, it is useful to consider an analogous framework from the study of distance processing in spatial cognition. Here, researchers use the concept of reference frames to describe the location of objects in physical space. Put simply, reference frames are means of describing the structure of the location of entities in space based on their relationships to other entities (Klatzky, 1998). In research on spatial perception, the two most commonly used reference frames are allocentric distance and egocentric distance. Allocentric distance describes the distances between each pair of objects within a given set. Egocentric distance identifies a particular object as the reference point origin and describes the distance between that object and each of the other objects within a set. Spatial reference frames have been a fruitful framework for studying navigation (Gramann et al., 2010), memory (Byrne et al., 2007), and computer vision (Henriques Vedaldi, 2018). Moreover, because allocentric and egocentric reference frames have a formal mathematical distinction, this framework has been useful in modeling brain activity for identifying shared and distinct systems for processing allocentric and egocentric distance in the underlying neurobiological mechanisms supporting each of these processes in humans (Wang, 2012; Zaehle et al., 2007) and non-human animals (Poulter et al., 2018; Rinaldi et al., 2020).

The utility of allocentric and egocentric reference frames can be directly applied to the study of social cognition where there is quantitative information about both the self and others within a group of people. Applying this framework to the domain of person knowledge, subjective ratings of the traits and characteristics of others and the self can serve as the basis of the coordinate systems where allocentric similarity describes the other-to-other differences and egocentric similarity describes self-to-other differences among a group of people. However, unlike spatial cognition, there is no objective ground-truth in person knowledge measures which are typically assessed through self-report ratings. Instead, researchers studying accuracy and agreement in interpersonal perception have made progress by aggregating responses among a set of raters to generate consensus-like judgments across participants (Kolar et al., 1996; McCrae Costa, 1987). Moreover, previous neuroimaging studies have demonstrated that consensus judgments of targets are better predictors of brain activity patterns than individual ratings (Knutson Genevsky, 2018) in some domains (e.g. affective responses during decision making) but less accurate in others (e.g. face-selective cortical areas; Kanwisher, 2017; Saygin et al., 2012). As such, it remains an open question whether an individual’s within-perceiver judgements or aggregated group-consensus responses are a better model of how reference frames of person knowledge are represented in the brains of individuals when they are considering the thoughts and behaviors of other people within their social groups.

The current study is designed to identify brain systems that represent information about other people based on either allocentric or egocentric reference frames of person knowledge within real-world social groups of well-acquainted others. Here, we successfully recruited 108 individuals from 20 independent groups of peers to make ratings about themselves and each other group using a round-robin design in which every person was both a perceiver and target for every other member of the group. Specifically, participants rated themselves and others on questions related to dimensions of warmth and competence from the stereotype content model (Fiske et al., 2002)—a robust and widely adopted theory of person perception of groups and individuals (Fiske, 2018) and their neural correlates (Thornton Mitchell, 2018). These ratings were then used to calculate within-perceiver allocentric and egocentric similarity measures among everyone in the group. Furthermore, group-consensus models of allocentric and egocentric similarity were calculated by averaging ratings across all members of the group. In a separate experimental session, the same participants were then scanned using functional magnetic resonance imaging (fMRI) while they completed a round-robin version of a standard trait-judgment task used to elicit mental representations of the self and others. Both subjective and consensus versions of each reference frame were then used as models of representational similarity that were compared against similarity of fMRI multivoxel brain responses patterns within the brains of participants when they were thinking of each other member of their group (see Fig. 1 for a general schematic of the approach). These analyses provide a means of identifying both shared and distinct neural systems in which representations of fellow group members are captured, simultaneously, by each reference frame of person knowledge during interpersonal perception, informing our understanding of reference frames for person perception and the brian systems that support these processes.

**Fig. 1.**
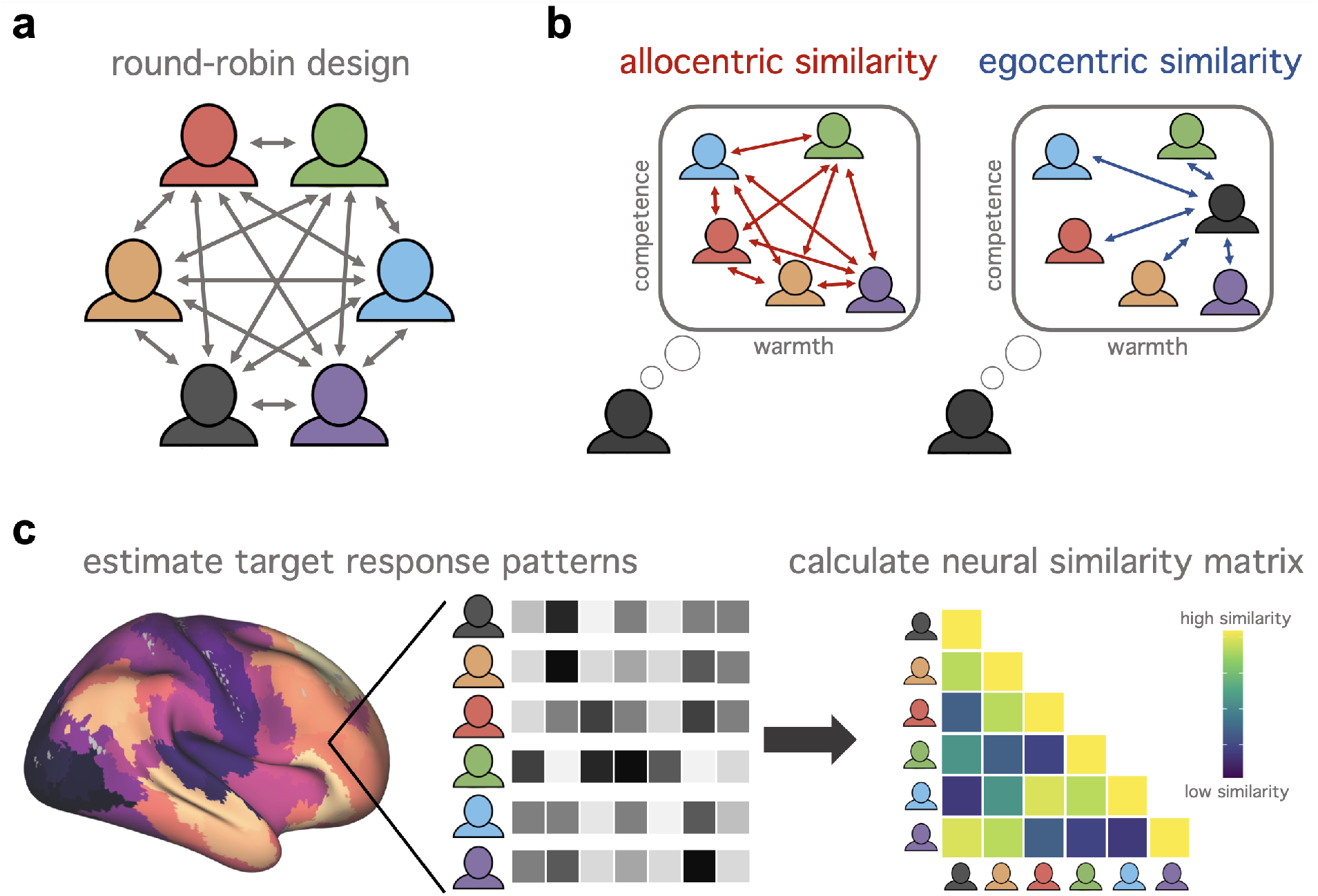
Design and analytic framework. A schematic of the approach used in the current study. A) schematic of the approach used in the current study. a, Participants were recruited from twenty independent social groups of six people in a round-robin design in which every participant is both a perceiver and target for every other member of their group. This round-robin design was used for both collecting behavioral ratings as well as measuring brain responses to each target. B) Allocentric similarity describes the distance in behavioral ratings among the members of each perceiver’s group. Egocentric similarity describes the distance in behavioral ratings between the perceiver and every other member of their group. Allocentric and egocentric reference frames were calculated using euclidean distances in behavioral ratings of measures of warmth and competence for ratings of both others and the self within each group. C) Brain activity for each target and the self were measured using functional magnetic resonance imaging within each participant. Multivoxel response patterns for each target and the self were then used to estimate the brain response similarity among each target within 600 cortical parcels. Behavioral models of allocentric and egocentric similarity were then compared to brain response similarity using a multilevel model representational similarity analysis to identify regions that were significantly associated with each reference frame within each participant.

## Methods

### Participants

A total of 120 right-handed participants were recruited from 20 independent, real-world social groups in the University of Oregon and local Eugene community, including student organizations, local business employees, and friend groups. Participants ranged in age from 18 to 51 (M = 23.5, SD = 7.2). A majority of participants identified as White or Caucasian (77.9%), 4.4% identified as Black or African American, 2.7% identified as Asian, 1.8 % identified as American Indian or Alaska Native, 1.8% identified as Native Hawaiian or Pacific Islander, 9.7% identified as more than one race, and 1.8% chose not to identify their race. A majority of participants identified as non-Hispanic (84.1%). Six people were recruited from each of the 20 groups and all participants within each group were familiar with one another. Across all groups, five participants failed to fulfill scheduled experimental session appointments, two subjects were excluded due to poor quality imaging data, and five subjects were excluded for missing or incomplete behavioral data. This left a total of N = 108 participants in the final sample used for all analyses. All participants were screened for magnetic resonance imaging contraindications and had normal or corrected-to-normal vision. Each participant in the study participated in two sessions. The first was a behavioral ratings session in which participants filled out a series of questionnaires for themselves and their group members. The second session consisted of a scanning session in which participants completed trait judgments for themselves and their peers while undergoing fMRI. Participants gave informed consent in both sessions in accordance with the guidelines set by the Internal Review Board at the University of Oregon and were compensated for their participation following each session of the study.

### Behavioral Rating Task

Participants were brought into the lab and asked to answer a series of questions about themselves and a set of familiar peers from within a small subset of their social network. Critically, we employed a round-robin design in which each participant in the study was both a perceiver and target stimulus for every other participant within their group (Figure 1a). All responses were recorded using PsychoPy stimulus presentation software (Peirce et al., 2019). During the behavioral trait ratings session, participants were presented with an array of scales (responses 1 to 5), each with the name of one of their peers or their own name listed above each. At the top of the screen, a single question was displayed, and participants were asked to give a rating corresponding to the impression of their peers or their evaluation of themselves on that given trait. Participants were required to make a rating for every peer and the self on the screen before being allowed to move on to the next question to ensure complete responses. Warmth and competence ratings from the twenty-one item stereotype content model (Fiske et al., 2002) were used as the measure of behavioral allocentric and egocentric reference frame calculation in the analysis (Figure 1b). In addition to measures from the stereotype content model, participants also completed ratings from other scales related to interpersonal closeness and mental health as part of a larger study on self-referential processing and interpersonal perception (see: Guthrie et al., 2022). The entire behavioral measurement data collection session lasted approximately one hour.

### Reference Frame Calculation

All reference frame models were calculated using item-wise responses of warmth and competence ratings of every target (including self ratings) per participant within each group. To this end, target ratings were flattened into one-dimensional response vectors that were then used to calculate the total euclidean distance among ratings for each target within each participant. Euclidean distance was used for behavioral measures because it captures both the direction and magnitude of the similarity among items, akin to spatial reference frames. For ease of interpretation, we use the term ‘similarity’ to describe the inverse of the euclidean distance metrics, such that smaller distances are equivalent to greater similarity. Unlike spatial reference frames, which have an objective criterion to establish distances among objects, reference frames of person knowledge are subjective and can be calculated according to judgements within the perceiver or via aggregated group consensus ratings. The round-robin design used in the current study allows for reference frame calculation both within-perceivers (i.e. within the person whose brain activity is being measured) and those averaged across fellow group members. This provides a means by which to know if group-consensus ratings provide a better model of a participant’s brain representation of target similarities than those based on the participant’s own subjective ratings, which may be biased or less accurate.

For the allocentric reference frame, within-perceiver similarity among targets was calculated based on that participant’s own ratings of the targets (i.e., the other members of their group) to generate a similarity matrix among all other members of the group. Group-consensus target similarity was calculated by averaging scores across items per target and recalculating reference frames within each group. Importantly, neither the perceiver’s trait ratings nor the individual target’s self-judgements were included in the aggregated results to avoid contaminating aggregated results with bias from either the target or perceiver.

Egocentric reference frames were calculated as the distance between ratings of the self and ratings of others. As with the allocentric reference frame, within-perceiver similarity was calculated using the ratings of the self from each participant against their ratings of all of their fellow group members. Unlike the allocentric reference frames, there are two different ways to calculate group-consensus responses for egocentric similarity. The first calculates the similarity of participant ratings of the self against group-aggregated responses of each target. The second calculates both group-aggregated ratings of the perceiver and group-aggregated ratings of the target. As with the allocentric reference frame, neither the perceiver’s trait ratings nor the individual target’s self-ratings were included in the group-aggregated calculations.

### Neuroimaging Data Acquisition

Magnetic resonance imaging was conducted with a Siemens 3T Skyra scanner using a 32-channel phased array coil. Structural images were acquired using a T1-weighted MP-RAGE protocol (175 sagittal slices; TR: 2500 ms; TE: 3.43 ms; flip angle: 7°; 1 mm isotropic voxels). Functional images were acquired using a T2*-weighted echo planar sequence (TR: 2000 ms; 72 axial slices; TE: 25 ms; flip angle: 90°; 2 mm isotropic voxels). For each participant, we collected five runs of the round-robin task (188 whole brain volumes per run). In order to correct for distortion due to B0 inhomogeneity we also acquired a field map (TR: 6390 ms; TE: 47.8 ms; effective echo spacing: 0.345 ms). The total length of time for the entire scanning session was approximately 75 minutes and each functional run was approximately 6 minutes long.

### Neuroimaging Trait Judgement Task

In a separate session, participants were brought back to the lab to complete the fMRI portion of the experiment. While in the scanner, participants were asked to complete a standard trait-judgment task widely used in the study of self and other processing (Chavez et al., 2017; Kelley et al., 2002; Mitchell et al., 2011). As with the behavioral ratings, we employed a round-robin design in which each participant in the study was both a perceiver and target stimulus for every other participant within their group. Participants were presented with a screen containing two words arranged vertically consisting of white text on a black background. For each trial, the top word displayed either “Self” or the name of one of the other five group members from the same group that the perceiver belonged to. Each perceiver indicated during the behavioral session what name they used most frequently when referring to each of their group members (e.g. “Robert” may have been changed to “Rob” or the nickname “Bean” for the fMRI portion). These names were then used in the fMRI portion to avoid confusion about which person each perceiver was making trait-judgements about for each trial. The bottom word displayed one of 60 valence-balanced trait adjectives (e.g. “Happy”, “Clumsy”, “Smart”; Anderson, 1968) for 2000 ms followed by 2000 ms of fixation and intermittent passive fixation trials (2000-12000 ms). Jittered trials were optimized using Opseq2 (Dale, 1999). Participants were asked to make either a “yes” or “no” response using a button box in their right hand as to whether or not the trait adjective described either themselves or one of their group members. All targets were presented in each run, and there were a total of twelve trials per target per run in which each target was presented with the same 12 trait-adjectives per run. The same targets were then used in each subsequent run but were then paired with a new set of 12 trait-adjectives in each one resulting in all targets being paired with all 60 trait-adjectives. Individual traits were only presented once per target randomly across all runs in the experiment. No two participants were presented with the same target/trait-adjective order across the experiment to account for potential order effects.

### Neuroimaging Preprocessing and Response Pattern Estimation

Functional imaging data for the fMRI task were preprocessed and voxel responses estimated using FSL (Smith et al., 2004). Data were processed using mean-based intensity normalization, high pass filtering (Gaussian-weighted least-squares straight line fitting, with sigma = 100s), and spatially smoothed with a 4mm FWHM Gaussian smoothing kernel. A multi-step normalization procedure was used to register results to standard space. First, functional data were corrected for spatial distortion using a field map unwarping before aligning functional data to each participant’s anatomical scan using boundary-based registration (Greve & Fischl, 2009) in conjunction with a linear registration with FSL’s FLIRT tool. These images were then warped to a 2mm MNI template using nonlinear registration with FSL’s FNIRT tool and a 10mm warp field. All first level task-based analyses were performed in native space before being warped into standard space for final analyses. Parameter estimates were calculated for each participant’s five group members in each of the five runs. These responses were then combined in a second-level within-subject fixed-effects analysis to produce parameter estimates for each of the five group members’ “target” conditions. Normalized (i.e. z-score) voxel responses for each condition were extracted from each parcel of the 600 regions in the Schaefer parcellation atlas (Schaefer et al., 2018). Consistent with previous studies using fMRI to estimate multivariate similarity patterns (Kriegeskorte, 2008; Thornton & Mitchell, 2018), we used Spearman rank correlation distance (e.g., 1 Spearman-ρ) to compute the dissimilarity between neural response patterns among each of the conditions (i.e. the self and each of the other people in the group) using tools from the nilearn package in Python (Abraham et al., 2014). These values were used as the estimates of neural similarity in each brain parcel that was then compared against the behavioral reference frame models (Figure 1c).

### Statistical Modeling

Behaviorally measured allocentric and egocentric reference frame similarity models were related to neural similarity measures at every parcel within each participant using a representational similarity analysis (Kriegeskorte, 2008). Allocentric similarity was modeled based on both within-perceiver and group-consensus ratings for targets. Egocentric similarity was modeled using within-perceiver ratings, group-consensus ratings for the target, and group-consensus ratings of both the perceiver and targets. For each reference frame, we fit a nested multilevel model using each behavioral similarity as a predictor of brain response similarity within nested random intercepts for each participant within each group using the lme4 package in R (Bates et al., 2015). Significance of each model was calculated using Satterthwaite estimated p-values for every parcel (Kuznetsova et al., 2017). Finally, whole-brain significance tests were corrected for multiple comparisons using false discovery rate (FDR) across all parcels.

## Results

### Allocentric Similarity

To increase the interpretability and transparency of the results, distributions of the behavioral measures and unthresholded statistical maps for the allocentric similarity analyses are shown in Figure 2a. There was a greater degree and wider distribution of allocentric distance values for within-perceiver effects (M = 4.82, SD = 1.89) than group-consensus for targets values (M = 3.45, SD = 1.11). Across the brain, within-perceiver models showed more consistent brain-behavior associations than group-consensus models. Unthresholded results were most pronounced in lateral prefrontal cortex and cortical midline structures from the default mode network.

**Fig. 2.**
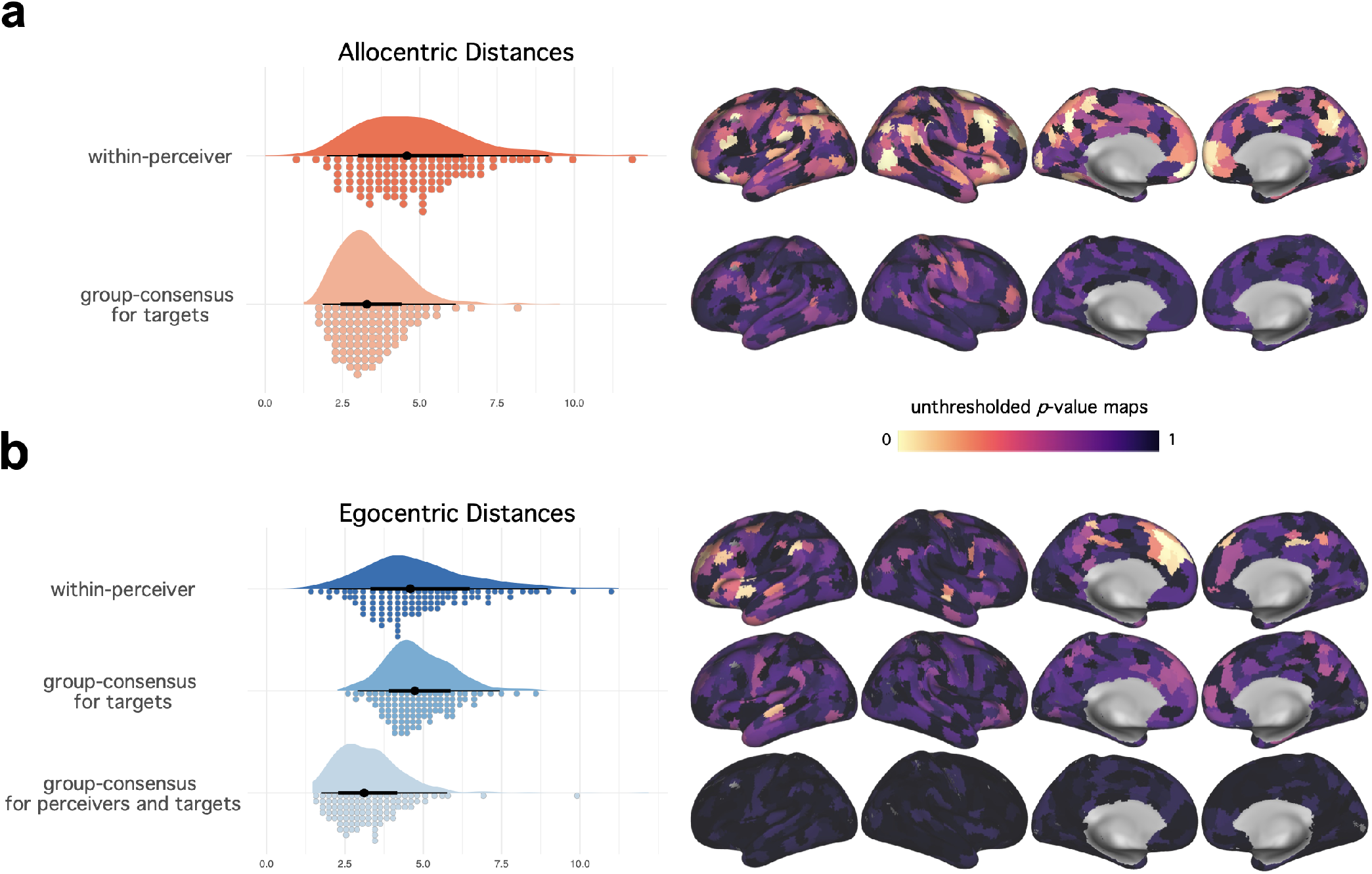
Unthresholded brain maps of reference frame representation. The distribution of behaviorally measured distance values for each reference frame and their associated unthresholded p-values from the fMRI representational similarity analysis. a, Allocentric distances among targets were calculated using both within-perceiver ratings and aggregated group-consensus ratings. Across the brain, within-perceiver ratings of allocentric similarity showed better model fit against neural similarity measures than the group-consensus ones, including within the default mode network and other areas implicated in social cognition. b, Egocentric distances were calculated using within-perceiver ratings, aggregated group-consensus ratings of only the targets against self-reported ratings from the perceiver, and aggregated group-consensus ratings of both the targets and the perceiver. Consistent with the allocentric results, the within-perceiver model of egocentric similarity showed better model fit against the neural similarity measures than either group-consensus models, particularly within prefrontal cortical regions.

### Egocentric Similarity

As with the allocentric reference frame results, behavioral distributions and unthresholded statistical maps from each of the egocentric similarity analyses are shown in Figure 2b. Unlike the allocentric reference frame results, distributions of ratings were similar between within-perceiver (M =4.82, SD = 1.78) and group-consensus for targets (M = 4.87, SD = 1.10). Group-consensus for targets and perceivers distance ratings were generally lower than the other two models (M = 3.30, SD = 1.23). Consistent with the results from the allocentric similarity analyses, only the within-perceiver effects showed brain-behavioral associations in multiple regions. In general, the results from the egocentric reference frame analyses were less distributed than the allocentric results and were mostly found in dorsal medial prefrontal cortical regions, as well as some lateral prefrontal areas.

### Non-overlapping Neural Representations of Person Knowledge Reference Frames

Results for the within-perceiver ratings after correcting for multiple comparisons for both allocentric and egocentric reference frames are shown in Figure 3. From these results, we found a positive relationship between multivoxel similarity patterns and behavioral allocentric similarity in parts of the brain previously implicated in social cognition, such as the posterior cingulate cortex, ventromedial prefrontal cortex, and lateral orbitofrontal cortex. There were also significant results found in areas previously implicated in spatial allocentric reference frame encoding (Zaehle et al., 2007), such as the inferior and superior frontal gyri and lateral occipital cortex. A table of all significant regions from the allocentric reference frame analysis can be found in Supplementary Table 1. There were no significant regions from the allocentric reference frame analysis that showed an inverse relationship to the behavioral similarity measures.

**Fig. 3.**
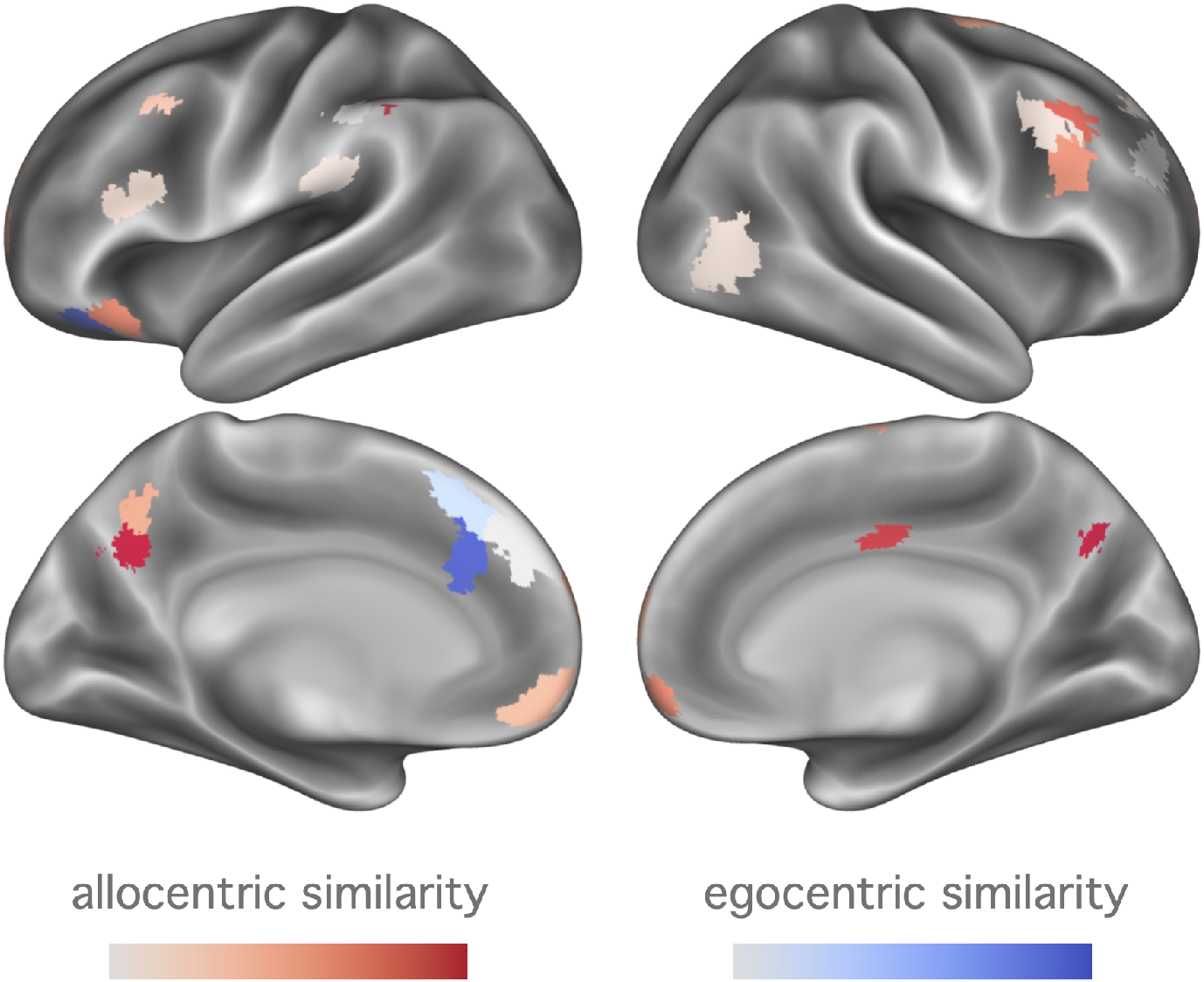
Non-overlapping regions reflecting reference frames of person knowledge. Thresholded FDR-corrected p-value maps of the significant areas from the within-perceiver allocentric similarity and egocentric similarity analyses. Brain and behavior allocentric similarity showed a positive relationship across a variety of regions involved in distance perception and social cognition, including the precuneus and ventral medial prefrontal cortex. Brain and behavior egocentric similarity showed an inverse relationship in regions of the the dorsal medial prefrontal cortex and anterior cingulate. There were no significant regions in which both allocentric similarity and egocentric similarity showed a significant relationship to the behavioral models of either reference frame.

From the egocentric reference frames results, we found a significant relationship between multivoxel similarity patterns and behavioral egocentric similarity, primarily within medial prefrontal regions including the dorsal medial prefrontal cortex and dorsal anterior cingulate cortex. A table of all significant regions from the egocentric reference frame analysis can be found in Supplementary Table 2. Importantly, all of the significant regions from the egocentric reference frame analysis showed an inverse relationship to the behavioral measures, such that greater egocentric similarity was related to less similar neural representations in these regions.

Finally, there were no significant results found after correcting for multiple comparisons for any of the group-consensus results across both reference frames. This suggests that subjective ratings from each participant were a better match to their own brain representations of group members than were ratings aggregated across fellow group members.

## Discussion

There are multiple ways humans can represent information about others that contribute to how we think of and interact with people in our social environments. In the current study, we adapted the notion of spatial allocentric and egocentric reference frames to measures of person knowledge to study how the brain represents information about otherto-other similarities and self-to-other similarities. To this end, we used a novel round-robin design for both behavioral and fMRI paradigms to relate models of person knowledge similarity to neural similarity during interpersonal perception within a variety of real-world social groups. Our results show that both allocentric similarity and egocentric similarity are related to measures of neural similarity, but that they did so separately with no overlap in the regions implicated for each reference frame. This suggests that reference frames for person knowledge are processed independently and concurrently, and not as two different modes of a single social cognitive system.

The results from the allocentric reference frame implicated a variety of brain regions involved in both social cognition and regions associated with spatial reference processing. Our results are consistent with a large body of work has implicated areas within the default mode network, including the ventral medial prefrontal context and posterior cingulate cortex, in processing information about other people (Denny et al., 2012; Wagner et al., 2012) and in representing similarities among famous individuals (Thornton Mitchell, 2018). Our results build on these previous findings to show the similarity of brain responses between real-world group members track with the similarity of major dimensions of person perception, namely assessments of warmth and competence among peers. Furthermore, our findings overlap with previous research that has also shown that brain regions such as the lateral occipital cortex, inferior frontal gyrus, and precuneus are involved in processing spatial allocentric distances (Galati et al., 2010; Zaehle et al., 2007). Together with these previous findings, our results are consistent with the idea that these regions which are engaged in spatio-temporal processes compute context general information that can be repurposed within social contexts (Parkinson Wheatley, 2015). Overall, the results of this study suggest that allocentric similarity is computed across broadly distributed cortical regions involved both in person perception and general distance processing.

The results from the egocentric reference frame analyses were different from the allocentric ones in a few important ways. First, brain regions related to egocentric reference frames were much less distributed, with results primarily located within the dorsal medial prefrontal cortex and anterior cingulate cortex, but also within the rostrolateral orbitofrontal cortex. These regions have been implicated in a variety of social cognitive tasks that involve social comparisons in close-others (Hughes Beer, 2012), self/other anchoring during mentalizing (Tamir Mitchell, 2010), and thinking of others with similar preferences (Mitchell et al., 2006) and were based on standard univariate responses magnitude metrics. Critically, our findings also showed that multivariate similarity of neural and behavioral measures for the egocentric similarity were inversely related to one another, such that the more similar a participant rated one of their peers, the more dissimilar their neural representation of that person was using our multivariate framework. This suggests that when people are reflecting on person knowledge about similar others, these brain systems may be engaging in processes that allow people to maintain a differentiation between others and the self. This interpretation is consistent with the view that a heightened or more distinguishable view of conspecifics from the self may allow a person to think about the mental states of others without confusing the source of those thoughts and feelings (de Waal, 2008), which may be critical for processes related to empathy in primates (de Waal Preston, 2017). Thus, at least when it comes to dissociating person knowledge of the self from similar others, these parts of the brain may be engaged not to underscore the shared aspects of those representations, but instead to ensure that we can maintain a cognitive separation of the self from group members with whom we have much in common.

The round-robin design used in the current study uniquely allowed us to measure each reference frame model based on subjective within-perceiver ratings of behavioral similarity as well as group-consensus ratings aggregated among fellow participants. This allowed us to test the possibility that group-consensus reference frames would predict neural similarity within each perceiver differentially than within-perciever ones. These analyses are critical because, unlike spatial reference frames, there is no objective way to define person knowledge similarity, and there may be potential issues with relating brain response similarity patterns to behavioral similarity metrics based on ratings that can be prone to self-serving or otherwise biased reporting. However, across both reference frames, we found no evidence that group-consensus behavioral ratings of any kind showed a superior relationship to the neural similarity measures. To the degree that within-perceiver ratings are prone to reporting biases, our results suggest that these biases are more consistent with how their brain is representing information about person knowledge than are those based on others’ ratings of the same target. Put differently, even if self-reports can be biased against an agreed upon social landscape, they may be reflecting honest interpretations of subjective similarities among people within our social groups and within our own brains.

## Conclusions

By integrating an interdisciplinary set of well-supported empirical models and methodological approaches, this study has theoretical implications for both cognitive neuroscience and social psychology. The results from our fMRI analyses indicate that independent regions known to be involved in various aspects of social cognition also process information about the differences among others as well as the differences between ourselves and others. Additionally, for the allocentric reference frame, we show that brain regions that are involved in more general cognitive distance processing also compute relevant information to person knowledge. For the egocentric reference frame, our results challenge assumptions about what information is being computed within brain regions when engaged in processing information concerning the differences between the self and others. Furthermore, our results inform social psychological theorizing by suggesting that our perception of other-to-other differences and self-to-other differences rely on independent social cognitive systems that may operate under different rules. To the degree that a social psychological theory posits that a manipulation or change in one reference frame implies a change in the other, our results suggest that this may not be the case and may offer a model for future investigations.

In summary, our results show that the brain independently processes both allocentric and egocentric reference frames of person knowledge among real-world groups of people. These findings contribute to our understanding of how the brain processes information about the self in social contexts which is central to our lives as richly sentient members of a fundamentally social species.

## ACKNOWLEDGEMENTS

Research funded by University of Oregon.

## OPEN SCIENCE PRACTICES

Data, materials, and code used to generate the findings in this study will be available prior to peer review via the Open Science Framework.

