## Supplementary Table 1 for "Person knowledge is independently encoded by allocentric and egocentric reference frames within separate brain systems"

*Significant FDR-Corrected Allocentric Reference Frame Parcels*

| Region | L/R | FDR p-value | Effect | SE | x | y | z |
|---|---|---|---|---|---|---|---|
| PCC | L | > .0001 | 0.0128 | 0.0023 | -6 | -64 | 34 |
| dLPFC | R | 0.0012 | 0.0104 | 0.0022 | 48 | 20 | 40 |
| lateral occipital cortex | R | 0.0024 | 0.007 | 0.0016 | 50 | -68 | 6 |
| supramarginal gyrus | L | 0.0032 | 0.0124 | 0.0029 | -54 | -46 | 54 |
| precuneus | R | 0.0032 | 0.0113 | 0.0027 | 6 | -68 | 38 |
| caudal lateral OFC | L | 0.0069 | 0.0098 | 0.0025 | -38 | 24 | -16 |
| lateral frontal pole | R | 0.0069 | 0.0087 | 0.0022 | 18 | 66 | 16 |
| vMPFC | R | 0.0086 | 0.0097 | 0.0025 | 8 | 64 | -12 |
| vMPFC | L | 0.0088 | 0.0074 | 0.0019 | -6 | 56 | -10 |
| dLPFC | L | 0.0183 | 0.0065 | 0.0018 | -46 | 12 | 40 |
| PCC | L | 0.0215 | 0.0084 | 0.0024 | -6 | -58 | 44 |
| lateral frontal pole | L | 0.0248 | 0.0088 | 0.0025 | -16 | 62 | 26 |
| middle frontal gyrus | R | 0.0248 | 0.0094 | 0.0027 | 44 | 18 | 28 |
| middle frontal gyrus | R | 0.0248 | 0.007 | 0.002 | 40 | 12 | 36 |
| dLPFC | R | 0.0278 | 0.0066 | 0.002 | 34 | 44 | 32 |
| motor cortex | L | 0.0303 | 0.004 | 0.0012 | -56 | -34 | 46 |
| somatosensory cortex | L | 0.0398 | 0.0047 | 0.0014 | -60 | -30 | 22 |
| motor cortex | R | 0.0398 | 0.0087 | 0.0027 | 20 | -6 | 70 |
| mid cingulate cortex | R | 0.0398 | 0.011 | 0.0034 | 6 | -10 | 38 |
| lateral inferior frontal gyrus | L | 0.0484 | 0.0051 | 0.0016 | -48 | 20 | 24 |
| lateral frontal pole | L | 0.0484 | 0.008 | 0.0025 | -24 | 64 | 12 |
| lateral frontal pole | R | 0.0484 | 0.007 | 0.0022 | 24 | 42 | 46 |

**Supplementary Table 2**

*Significant FDR-corrected Egocentric Reference Frame Parcels*

| Region | L/R | FDR p-value | Effect | SE | x | y | z |
|---|---|---|---|---|---|---|---|
| rostral lateral OFC | L | 0.0172 | -0.0107 | 0.0025 | -4 | 28 | 48 |
| dMPFC | L | 0.0444 | -0.0096 | 0.0025 | -4 | 48 | 36 |
| medial superior frontal cortex | L | 0.0468 | -0.0144 | 0.004 | -8 | 28 | 30 |
| dACC | L | 0.0468 | -0.0142 | 0.0039 | -36 | 38 | -12 |

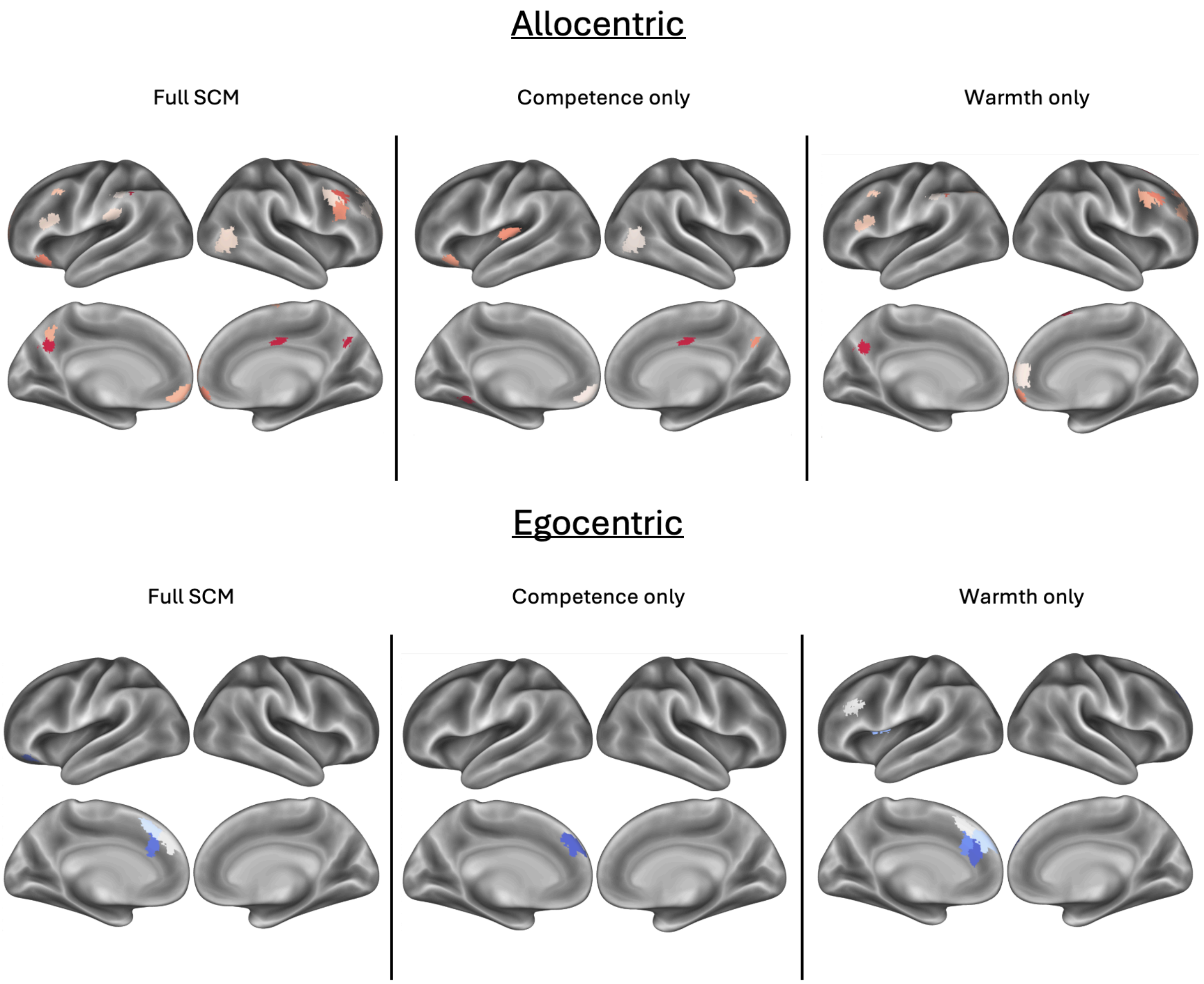


**Supplementary Figure 1.** Results from reference frames based on warmth and competence dimensions separately compared to the full stereotype content model (SCM) of both dimensions together. All results shown are thresholded FDR-corrected *p*-value maps of the significant areas from the within-perceiver allocentric similarity and egocentric similarity analyses.
